## Supplemental material for "The influence of nucleus accumbens shell D1 and D2 neurons on outcome-specific Pavlovian instrumental transfer"

### Supplemental Figures

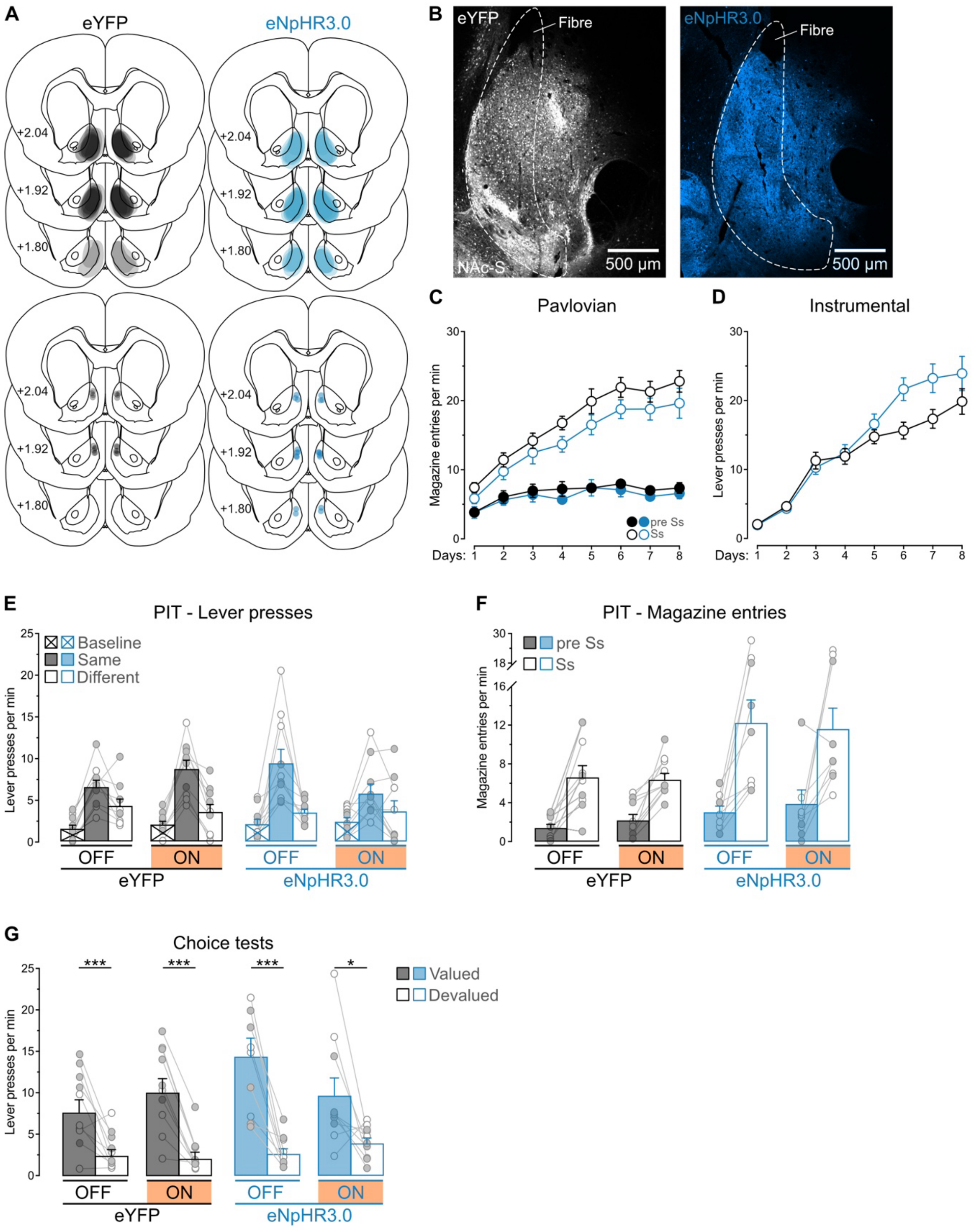


**Supplemental Figure S1 – Histological and behavioral data related to Figure 3**

**A.** Minimal (light gray for eYFP group and light blue for eNpHR3.0 group) and maximal (darker gray for eYFP group and darker blue for eNpHR3.0 group) extent of the NAc-S viral infection in D1-Cre rats. Location of fiber-optic cannulas in the NAc-S (gray for eYFP group and blue for eNpHR3.0 group) are also represented. Distances are indicated in mm from bregma. **B**. Micrographs showing viral expression in NAc-S D1 SPNs in the eYFP group (left) and eNpHR3.0 group (right). **C**. During Pavlovian conditioning, magazine entry rates were similar across group (Group – eYFP vs. eNpHR3.0: $p = 0.13$) and increased across days (Day: $F_{1,18} = 88.24$, $p < 0.001$, $\eta^2 = 0.83$) regardless of group (Group x Day: $p = 0.45$). The rates were higher in the presence of the stimuli than in their absence (Period, S vs. pre: $F_{1,18} = 259.04$, $p < 0.001$, $\eta^2 = 0.94$) irrespective of group (Group x Period: $p = 0.12$). The discrimination between the two periods increased as training progressed (Day x Period: $F_{1,18} = 66.84$, $p < 0.001$, $\eta^2 = 0.79$) regardless of group (Group x Day x Period: $p = 0.72$). **D**. During instrumental conditioning, lever press rates were similar across group (Group: $p = 0.16$) and increased as training progressed (Day: $F_{1,18} = 190.83$, $p < 0.001$, $\eta^2 = 0.91$). The increase was more pronounced in the eNpHR3.0 group than in the eYFP group (Group x Day: $F_{1,18} = 5.40$, $p < 0.05$, $\eta^2 = 0.23$). **E**. Raw data for lever presses during the PIT test. **F**. During the PIT test, magazine entry rates were higher in the eNpHR3.0 group than the eYFP group (Group: $F_{1,18} = 7.84$, $p < 0.05$, $\eta^2 = 0.30$). The rates remained unaffected by LED activation (LED – ON vs. OFF, $p = 0.16$) regardless of group (Group x LED: $p = 0.89$). The rates were higher in the presence of the stimuli than in their absence (Period: $F_{1,18} = 26.58$, $p < 0.001$, $\eta^2 = 0.60$) irrespective of group (Group x Period: $p = 0.12$). The discrimination between the two periods was unaffected by LED activation (Period x LED: $p = 0.15$) irrespective of group (Group x Period x LED: $p = 0.78$). **G**. During the choice test, lever press rates were similar across groups (Group: $p = 0.15$) and were higher on the lever earning the valued outcome (Lever – Valued vs. Devalued: $F_{1,18} = 49.03$, $p < 0.001$, $\eta^2 = 0.74$) irrespective of group (Group x Lever: $p = 0.33$). Lever press rates were unaffected by LED activation (LED: $p = 0.61$) irrespective of group (Group x LED: $p = 0.06$). The difference between rates on the valued and devalued lever was left intact by LED activation (Lever x LED: $p = 0.26$) but this depended on the group considered (Group x Lever x LED: $F_{1,18} = 10.04$, $p < 0.01$, $\eta^2 = 0.36$). Nevertheless, simple effect analyses revealed that lever press rates were higher on the valued lever compared to the devalued lever in both groups whether the LED was OFF or ON (eYFP-OFF: $F_{1,9} = 13.29$, $p<0.01$, $\eta^2 = 0.57$; eYFP-ON: $F_{1,9} = 47.45$, $p < 0.001$, $\eta^2 = 0.84$; eNpHR3.0-OFF: $F_{1,9} = 31.68$, $p < 0.001$, $\eta^2 = 0.78$; eNpHR3.0-ON: $F_{1,9} = 6.36$, $p < 0.05$, $\eta^2 = 0.41$). Data are shown as mean ± SEM. Panels E-G include individual data points for female (filled circle) and male (open circle) rats. Asterisks denote significant effect (***p<0.001; *p<0.05), n.s., nonsignificant.

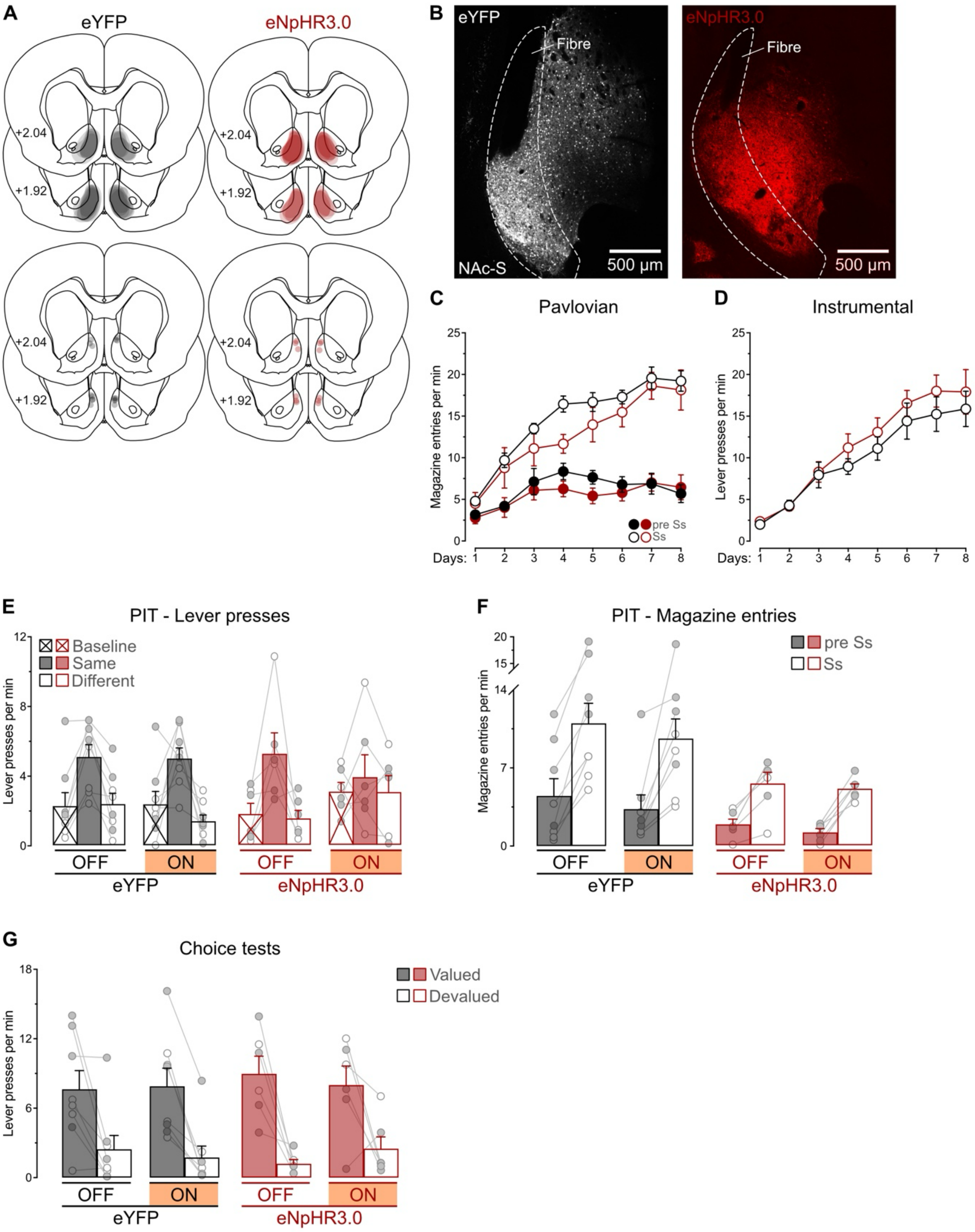


**Supplemental Figure S2 – Histological and behavioral data related to Figure 4**
**A**. Minimal (light gray for eYFP group and light red for eNpHR3.0 group) and maximal (darker gray for eYFP group and darker red for eNpHR3.0 group) extent of the NAc-S viral infection in A2a-Cre rats. Location of fiber-optic cannulas in the NAc-S (gray for eYFP group and red for eNpHR3.0 group) are also represented. Distances are indicated in mm from bregma. **B**. Micrographs showing viral expression in NAc-S D2 SPNs in the eYFP group (left) and eNpHR3.0 group (right). **C**. During Pavlovian conditioning, magazine entry

rates were similar across group (Group: $p = 0.26$) and increased across days (Day: $F_{1,12} = 89.63$, $p < 0.001$, $\eta^2 = 0.88$) regardless of group (Group x Day: $p = 0.85$). The rates were higher in the presence of the stimuli than in their absence (Period: $F_{1,12} = 160.90$, $p < 0.001$, $\eta^2 = 0.93$) irrespective of group (Group x Period: $p = 0.39$). The discrimination between the two periods increased as training progressed (Day x Period: $F_{1,12} = 74.66$, $p < 0.001$, $\eta^2 = 0.86$) regardless of group (Group x Day x Period: $p = 0.70$). **D**. During instrumental conditioning, lever press rates were similar across group (Group: $p = 0.45$) and increased as training progressed (Day: $F_{1,12} = 116.85$, $p < 0.001$, $\eta^2 = 0.91$) irrespective of group (Group x Day: $p = 0.39$). **E**. Raw data for lever presses during the PIT test. **F**. During the PIT test, magazine entry rates were similar across groups (Group: $p = 0.053$). The rates remained unaffected by LED activation (LED: $p = 0.055$) regardless of group (Group x LED: $p = 0.44$). The rates were higher in presence of the stimuli than in their absence (Period: $F_{1,12} = 50.37$, $p < 0.001$, $\eta^2 = 0.81$) irrespective of group (Group x Period: $p = 0.09$). The discrimination between the two periods was unaffected by LED activation (Period x LED: $p = 0.89$) irrespective of group (Group x Period x LED: $p = 0.75$). **G**. During the choice test, lever press rates were similar across groups (Group: $p = 0.86$) and were higher on the lever earning the valued outcome (Lever: $F_{1,12} = 66.95$, $p < 0.001$, $\eta^2 = 0.85$) irrespective of group (Group x Lever: $p = 0.54$). Lever press rates were unaffected by LED activation (LED: $p = 0.98$) irrespective of group (Group x LED: $p = 0.85$). The difference between rates on the valued and devalued lever was left intact by LED activation (Lever x LED: $p = 0.71$) regardless of group (Group x Lever x LED: $p = 0.36$). Data are shown as mean ± SEM. Panels E-G include individual data points for female (filled circle) and male (open circle) rats.

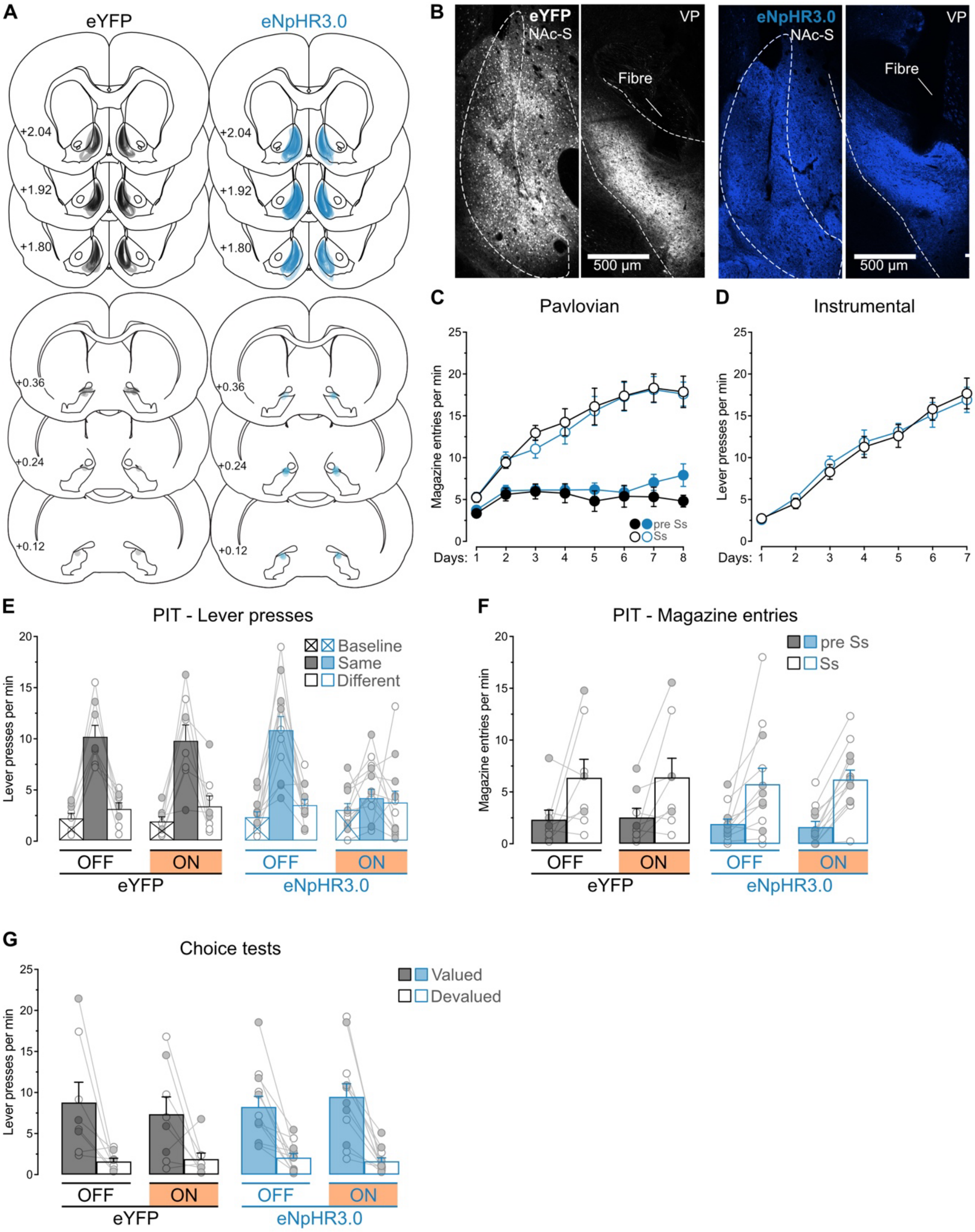


**Supplemental Figure S3 – Histological and behavioral data related to Figure 5**

**A**. Minimal (light gray for eYFP group and light blue for eNpHR3.0 group) and maximal (darker gray for eYFP group and darker blue for eNpHR3.0 group) extent of the NAc-S viral infection in D1-Cre rats. Location of fiber-optic cannulas in the VP (gray for eYFP group and blue for eNpHR3.0 group) are also represented. Distances are indicated in mm from bregma. **B**. Micrographs showing viral expression in NAc-S D1 SPNs and their terminals in the VP for the eYFP group (left) and eNpHR3.0 group (right). **C**. During Pavlovian conditioning, magazine entry rates were similar across group (Group: $p = 0.79$) and increased across days (Day: $F_{1,18}$ = 83.14, $p < 0.001$, $\eta^2$ = 0.82) regardless of group (Group x Day: p = 0.50). The rates were higher in the presence of the stimuli than in their absence (Period: $F_{1,18}$ = 129.21, $p < 0.001$, $\eta^2$ = 0.88) irrespective of group (Group x Period: p = 0.31). The discrimination between the two periods increased as training progressed (Day x Period: $F_{1,18}$ = 48.18, $p < 0.001$, $\eta^2$ = 0.73) regardless of group (Group x Day x Period: p = 0.49). **D**. During instrumental conditioning, lever press rates were similar across groups (Group: p = 0.90) and increased as training progressed (Day: $F_{1,18}$ = 171.16, $p < 0.001$, $\eta^2$ = 0.90) irrespective of group (Group x Day: p = 0.65). **E**. Raw data for lever presses during the PIT test. **F**. During the PIT test, magazine entry rates were similar across group (Group: p = 0.72). The rates remained unaffected by LED activation (LED: p = 0.77) regardless of group (Group x LED: p = 0.96). The rates were higher in the presence of the stimuli than in their absence (Period: $F_{1,18}$ = 20.80, $p < 0.001$, $\eta^2$ = 0.54) irrespective of group (Group x Period: p = 0.86). The discrimination between the two period was unaffected by LED activation (Period x LED: p = 0.73) irrespective of group (Group x Period x LED: p = 0.56). **G**. During the choice test, lever press rates were similar across group (Group: p = 0.72) and were higher on the lever earning the valued outcome (Lever: $F_{1,18}$ = 36.12, $p < 0.001$, $\eta^2$ = 0.67) irrespective of group (Group x Lever: p = 0.97). Lever press rates were unaffected by LED activation (LED: p = 0.89) irrespective of group (Group x LED: p = 0.45). The difference between rates on the valued and devalued lever was left intact by LED activation (Lever x LED: p = 0.99) regardless of group (Group x Lever x LED: p = 0.27). Data are shown as mean ± SEM. Panels E-G include individual data points for female (filled circle) and male (open circle) rats.

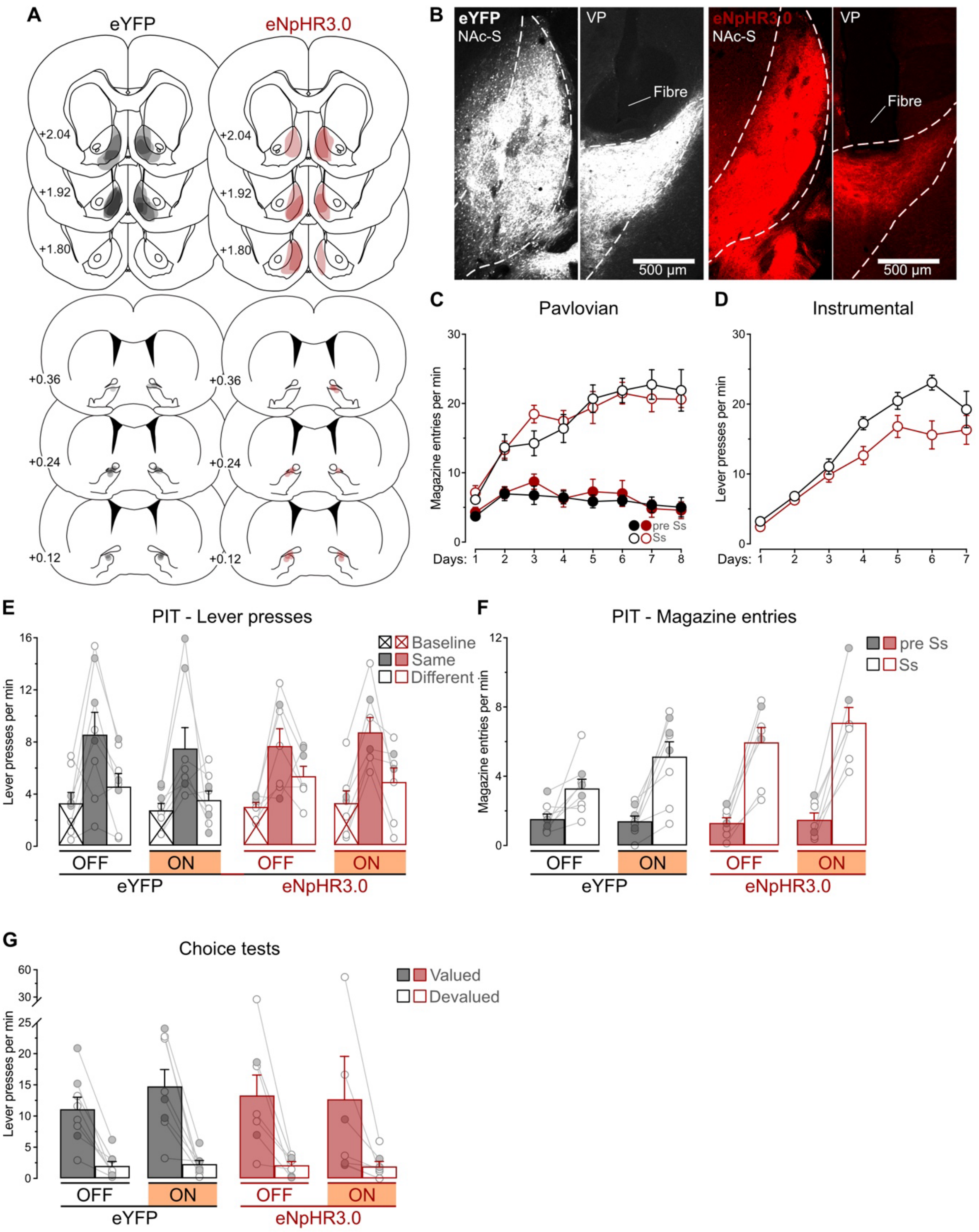


**Supplemental Figure S4 – Histological and behavioral data related to Figure 6**

**A**. Minimal (light gray for eYFP group and light red for eNpHR3.0 group) and maximal (darker gray for eYFP group and darker red for eNpHR3.0 group) extent of the NAc-S viral infection in A2a-Cre rats. Location of fiber-optic cannulas in the VP (gray for eYFP group and red for eNpHR3.0 group) are also represented. Distances are indicated in mm from bregma. **B**. Micrographs showing viral expression in NAc-S D2 SPNs and their terminals in the VP for the eYFP group (left) and eNpHR3.0 group (right). **C**. During Pavlovian conditioning, magazine entry rates were similar across group (Group: $p$ = 0.83) and increased across days (Day: $F_{1,13}$ = 23.82, $p < 0.001$, $\eta^2 = 0.65$) regardless of group (Group x Day: p = 0.42). The rates were higher in the presence of the stimuli than in their absence (Period: $F_{1,13}$ = 767.22, $p < 0.001$, $\eta^2 = 0.98$) irrespective of group (Group x Period: p = 0.64). The discrimination between the two periods increased as training progressed (Day x Period: $F_{1,13}$ = 106.62, $p < 0.001$, $\eta^2 = 0.89$) regardless of group (Group x Day x Period: p = 0.40). **D**. During instrumental conditioning, lever press rates were higher in the eYFP group than the eNpHR3.0 group (Group: $F_{1,13}$ = 6.18, $p < 0.03$, $\eta^2 = 0.32$). The rates increased as training progressed (Day: $F_{1,13}$ = 155.04, $p < 0.001$, $\eta^2 = 0.92$) irrespective of group (Group x Day: p = 0.10). **E**. Raw data for lever presses during the PIT test. **F**. During the PIT test, magazine entry rates were similar across groups (Group: p = 0.09). Larger rates were observed with LED activation (LED: $F_{1,13}$ = 6.26, $p < 0.05$, $\eta^2 = 0.32$) but this was true regardless of group (Group x LED: p = 0.74). The rates were higher in the presence of the stimuli than in their absence (Period: $F_{1,13}$ = 76.18, $p < 0.001$, $\eta^2 = 0.86$), especially in the eNpHR3.0 group compared to the eYFP group (Group x Period: $F_{1,13}$ = 6.97, $p < 0.05$, $\eta^2 = 0.35$). The discrimination between the two period was increased by LED activation (Period x LED: $F_{1,13}$= 11.17, $p < 0.01$, $\eta^2 = 0.46$) but this was true regardless of group (Group x Period x LED: p = 0.26). **G**. During the choice test, lever press rates were similar across groups (Group: p = 0.99) and were higher on the lever earning the valued outcome (Lever: $F_{1,13}$ = 27.04, $p < 0.001$, $\eta^2 = 0.68$) irrespective of group (Group x Lever: p = 0.97). Lever press rates were unaffected by LED activation (LED: p = 0.67) irrespective of group (Group x LED: p = 0.51). The difference between rates on the valued and devalued lever was left intact by LED activation (Lever x LED: p = 0.62) regardless of group (Group x Lever x LED: p = 0.53). Data are shown as mean ± SEM. Panels E-G include individual data points for female (filled circle) and male (open circle) rats.

### Supplemental discussion

#### Magazine entry rates during the PIT tests

NAc-S D1-SPNs silencing enhanced magazine entry rates during the outcome-specific PIT test (Supplemental Figure S1F). During this test, stimuli elicit both magazine entries and lever presses, with these response types competing for behavioral control (Lovibond, 1981). Manipulations that reduce one response type might therefore be expected to enhance the other. Since NAc-S D1-SPNs silencing impaired outcome-specific PIT and reduced lever press responding (Figure 3C and Supplemental Figure S1E), it would seem reasonable to conclude that this reduction resulted in increased magazine entry rates. However, this increase was also observed during LED OFF trials in eNpHR3.0 rats (Supplemental Figure S1F), when lever press performance was comparable to control eYFP rats (Figure 3C and Supplemental Figure S1E). Therefore, response competition cannot explain the enhanced magazine entry rates. The results from NAc-S D2-SPNs silencing further rule out this possibility. While D2-SPNs silencing impaired outcome-specific PIT (Figure 4C and Supplemental Figure S2E), it had no significant effect on magazine entry rates during PIT, unlike D1-SPNs silencing. If anything, there was a trend toward reduced magazine entries with D2-SPNs silencing, but this reduction was also present during LED OFF conditions. Notably, a similar trend occurred during Pavlovian training, when lever press manipulanda were unavailable and no silencing had been implemented (Supplemental Figure 2C). Overall, the present data do not provide a clear explanation for the enhanced magazine entry rates observed in eNpHR3.0 rats during the PIT test. Future studies using pure Pavlovian tasks without a competing instrumental response may be necessary to establish whether NAc-S SPNs modulate magazine entry performance.

#### Choice tests under NAc-S D1-SPNs silencing

The choice test data presented in Supplemental Figure 1G revealed a significant Group × Lever × LED interaction. Follow-up analyses demonstrated that all groups retained the capacity to select between actions based on outcome value. However, the significant interaction suggested that NAc-S D1-SPNs silencing marginally attenuated this capacity. Caution is warranted before interpreting this finding as evidence for a role of NAc-S D1-SPNs in value-based decision-making. Several studies have demonstrated that NAc-S manipulations typically preserve such choice behavior (Corbit et al., 2001; Corbit & Balleine, 2011; Laurent et al., 2012). Furthermore, previous work showed that NAc-S D1 receptor blockade impairs outcome-specific PIT while leaving value-based choice unaffected (Laurent et al., 2014). We therefore favor an alternative explanation for the observed marginal reduction. As shown in Supplemental Figure 1A, viral transduction extended slightly into the nucleus accumbens core (NAc-C), a region critical for value-based decision-making (Corbit et al., 2001; Corbit & Balleine, 2011; Laurent et al., 2012; Parkes et al., 2015). The marginal impairment may therefore reflect inadvertent silencing of a small number of NAc-C D1-SPNs rather than a functional contribution from NAc-S D1-SPNs. Future studies specifically targeting a larger population of NAc-C D1-SPNs populations would help clarify this possibility.
